# Reconstituting the genus *Mycobacterium*

**DOI:** 10.1101/2021.03.11.434933

**Authors:** Conor J Meehan, Roman A. Barco, Yong-Hwee E Loh, Sari Cogneau, Leen Rigouts

## Abstract

The definition of a genus has wide ranging implications both in terms of binomial species names and also evolutionary relationships. In recent years, the definition of the genus *Mycobacterium* has been debated due to the proposed split of this genus into five new genera (*Mycolicibacterium, Mycolicibacter, Mycolicibacillus, Mycobacteroides* and an emended *Mycobacterium*). Since this group of species contains many important obligate and opportunistic pathogens, it is important that any renaming of species is does not cause confusion in clinical treatment as outlined by the *nomen periculosum* rule (56a) of the Prokaryotic Code.

In this study, we evaluated the proposed and original genus boundaries for the mycobacteria, to determine if the split into five genera was warranted. By combining multiple approaches for defining genus boundaries (16S rRNA gene similarity, amino acid identity index, average nucleotide identity, alignment fraction and percentage of conserved proteins) we show that the original genus *Mycobacterium* is strongly supported over the proposed five-way split. Thus, we propose that the original genus label be reapplied to all species within this group, with the proposed five genera used as sub-genus complex names.

## Introduction

The genus *Mycobacterium* was first named in 1896 by Lehmann and Neumann, based primarily on the features of the type strain *Mycobacterium tuberculosis* (Lehmann and Neumann, 1896). Such phenotypic characteristics include the presence of mycolic acids in the cell wall, aerobic growth and bacillary cell shape. Over the years, the taxonomic definition has been reinforced by 16S and phylogenomic analyses (Tortoli *et al*., 2017). The genus contains over 170 named species, including major human pathogens such as *M. tuberculosis* and *M. leprae* (Tortoli *et al*., 2017, 2019). It also encompasses the non-tuberculous mycobacteria (NTMs), many of which are major opportunistic pathogens such as *M. abscessus* and *M. avium*. The vast majority of mycobacterial species are environmental and can be found in a wide array of niches.

Based on phenotypic and phylogenomic data, the genus is often split first into rapid and slow growers and then further split into specific complexes or groups (e.g. *M. tuberculosis* complex or *M. avium* complex)(Fedrizzi *et al*., 2017). In 2018, the genus was split into four new genera: *Mycolicibacterium, Mycolicibacter, Mycolicibacillus, Mycobacteroides* and an emended *Mycobacterium* (Gupta, Lo and Son, 2018). This was done first using a whole genome sequenced(WGS)-based phylogeny, which revealed five major groups within the original genus, corresponding loosely to many of the previously described complexes (Fedrizzi *et al*., 2017). These groupings were then defined as genera based upon average amino acid identity (AAI), conserved signature indels (CSIs) and conserved signature proteins (CSPs).

Such taxonomic changes have far reaching consequences for genera that contain a considerable number of clinically important pathogens. Currently, the original genus name *Mycobacterium* serves as a synonym for these five genera, adding to the confusion around species naming. The recent splits potentially cause much confusion for clinical treatment of mycobacterial diseases as some major opportunistic pathogens such as *M. abscessus* have been renamed. Indeed, the commonly used term non-tuberculous mycobacteria is called into question as this now includes many species no longer defined as mycobacteria. The renaming also means that the instructions for use (IFUs) for NTM diagnostics, such as the Hain GenoType system (Hain LifeSciences, Germany) or similar, may need to be changed as the species listed are not the current validly published species names, which could cause additional confusion for clinicians. Thus, there is a need to ensure this split is strongly supported before clinical guidelines and IFUs are updated.

Despite being a widely used taxonomic rank, the definition of genus is somewhat elusive. Generally, it is defined as the taxonomic level above species and below family, without concrete methodology to circumscribe such a grouping. Phenotypic definitions usually include a combination of attributable features such as Gram staining or spore formation and biochemical test results such as catalase activity or amino acid degradation (Gupta, 2015). However, a unified genotypic approach has never been applied across several genera.

Early attempts to delimit genera included the use of the 16S rRNA gene identity cut-off of 94-95% (Yarza *et al*., 2008, 2014), and amino acid identity (AAI) score of ≥65% (Konstantinidis, Rosselló-Móra and Amann, 2017). These options have now been expanded and include three genome-based methods. The Relative Evolutionary Distance (RED) is the basis of the taxonomic structure underpinning the Genome Taxonomy Database (GTDB) (Parks *et al*., 2018). This method uses a conserved set of proteins to build a phylogeny and assign taxonomic ranks based on normalised evolutionary distances. Within this approach, the original genus *Mycobacterium* is preserved as a single genus, with no evidence for sub-splitting above the level of species and is listed as such in the GTDB (*GTDB - Mycobacterium tree*, 2021). A second method, termed Percentage of Conserved Proteins (POCP) uses a BLAST-based approach to define homologs between species (Qin *et al*., 2014). Two species belong to the same genus if 50% of their proteins are shared. This allows for amino acid divergence and protein evolution over long time spans, as is common for members of the same genus, while still ensuring core functionality and characteristics are shared. This method has gained popularity for defining a genus and has been used many times before (Adeolu *et al*., 2016; Lopes-Santos *et al*., 2017; Nicholson *et al*., 2018), but still has some issues as it relies on the 16S rRNA gene as a reference point. A third method uses a combination of the genome alignment fraction (AF) and the ANI to define genera boundaries (Barco *et al*., 2020). This method is not directly dependent on the 16S rRNA gene, unlike POCP.

In this work, we apply the POCP and AF/ANI methods to the original *Mycobacterium* genus and the newly proposed five genera of Gupta (Gupta, Lo and Son, 2018). We show that the original genus fits the POCP definition of genus while the new genera have overlapping boundaries between genera, making their definitions unsupported. The AF/ANI method also shows that splitting the genus into five genera is unfounded, although the designation of *Mycobacteroides* as a separate genus warrants further investigation. We thus propose that the five newly created genera should be reconstituted into a single genus, named *Mycobacterium*.

## Methods

The original designation of *Mycobacterium* before the genus was split into 5 genera (Gupta, Lo and Son, 2018) will be used here for clarity of the dataset under investigation. When referring to the new *Mycobacterium* genus this will be explicitly stated as *Mycobacterium* (Gupta).

### Dataset

All genome sequences used in previous phylogenomic studies of the genus *Mycobacterium* (Fedrizzi *et al*., 2017; Tortoli *et al*., 2017) were used here, covering the species re-designated in (Gupta, Lo and Son, 2018) and the corrigendum (Gupta, Lo and Son, 2019).

An outgroup was required to understand the boundary of genus delimitation for the proposed genus. To this end, all genomes in the GTDB (Parks *et al*., 2018, 2020) assigned to *Corynebacteriaceae*, the same family as the genus *Mycobacterium*, were retrieved from NCBI on the 31^st^ of January 2019.

A list of all genomes, their designation in the new genera, and their accession numbers are included in Supplementary Table 1.

### Genome quality control and annotation

Since annotation of the proteins for each species is vital for the proposed analyses, we wanted to ensure that a uniform approach was used for the detection and assigning of open reading frames. The program Prokka v1.13.7 (Seemann, 2014) was used to undertake genome annotation using the genomic sequence file retrieved from NCBI for each genome.

The program CheckM (Parks *et al*., 2015) was employed to ensure that only genomes of good quality were used for this study. The genomic sequence file (.fna) for each genome was input to CheckM. Those with an estimated genome completeness lower than 80% (indicating incomplete sequencing (Parks *et al*., 2015)) were removed from further analysis.

### 16S rRNA gene similarity and Average amino-acid identity (AAI) score estimation

The percentage similarity between the 16S sequences of each strain was calculated in a pairwise manner on the 16S sequences as annotated by Prokka using BLASTn, as implemented in BLAST+ v2.5 (Altschul *et al*., 1990).

The AAI between the proteomes of two strains is an estimate of their molecular relatedness using the amount of shared amino acids in their protein complements as a marker of divergence (Konstantinidis and Tiedje, 2005; Konstantinidis, Rosselló-Móra and Amann, 2017). Although this has been shown to be limited for delineating adjacent taxonomic ranks (e.g. species from genus and genus from family) it is still often used for this task. We used CompareM (Parks, 2014) to compute the AAI between all genomes in our dataset (i.e. the family *Corynebacteriaceae*). The package ggplot2 (Warnes, Bolker and Lumley, 2012) implemented in R v3.5.1 (R Core Team, 2017) was used to construct histograms of these pairwise AAI scores.

### Percentage of Conserved Proteins (POCP)

Due to the drawbacks of the AAI method for delineating genera, Qin *et al*. implemented a method based on shared protein content (i.e. presence/absence of proteins in both strains) as a measure of relatedness (Qin *et al*., 2014). Briefly, the POCP is calculated using the formula [(C1+C2)/(T1+T2)]*100% where C1 is the number of proteins in strain 1 also present in strain 2 and T1 is the total protein count of strain 1. C2/T2 are the equivalent numbers for strain 2.

This method was applied to our data by first undertaking reciprocal BLASTp searches using BLAST+ v2.5 with an e-value cut-off of 1e^-5^. A python 2.7 script was then used to filter these results, retaining only those with >40% similarity. These similarity and e-value cut-offs were selected as they are the settings used by Qin *et al*. A second python script was used to calculate the POCP. These scripts can be found at https://github.com/conmeehan/gentax. The resulting spread of POCP scores was visualised as a histogram using the ggplots2 (Warnes, Bolker and Lumley, 2012) package in R.

### AF and ANI

Alignment fraction (AF) and average nucleotide identity (ANI) values were obtained by using ANIcalculator 2014-127, version 1.0 (https://ani.jgi.doe.gov/html/download.php?)(Varghese et al., 2015). A custom perl script was written to automate submission of sequence comparisons to ANIcalculator; this can be found at https://github.com/eddieloh-usc/run_ANIcalculator. The genus demarcation approach that uses AF and ANI determines if a specific set of species constitutes a genus by using the type species of such genus as a stable reference for pairwise-comparisons, which include genera within the same taxonomic order or family (Barco *et al*., 2020). Since the genomic content of type strains is used to delineate genera, this makes the approach independent of a benchmarking ribosomal gene (e.g. 16S rRNA), which could introduce biases, as seen with other approaches. The spread of AF/ANI scores and the genus demarcation boundaries were visualised as a scatterplot using ggplots2.

## Results

A total of 366 genomes were included in this study: 148 genomes from the original *Mycobacterium* genus with the remaining 218 genomes from other genera in the family *Corynebacteriaceae*. Of these genomes, 13 were found to be incomplete by CheckM and were removed: nine *Mycobacterium*, two *Gordonia* and two *Rhodococcus*. This resulted in a dataset of 340 genomes, of which 139 were *Mycobacterium*, representing 123 *Mycobacterium* species based on the updated naming conventions set out by Tortoli *et al*. (Tortoli *et al*., 2019). The remaining 16 “mycobacterial” genomes represent subspecies that used to be designated as separate species (Tortoli *et al*., 2019), and four major *M. tuberculosis* lineages, besides lineage 4 (H37Rv reference strain) (Gagneux, 2018).

### 16S rRNA gene similarity and AAI support the original *Mycobacterium* designation

Although 16S rRNA gene similarity and AAI are not recommended for genus delineation (Konstantinidis and Tiedje, 2005; Qin *et al*., 2014), they have been used for such in the past. Generally, a cut-off of >94% is supportive of species belonging to the same genus. 16S similarity scores were compared within each genus (intra-genus) and between members of each specific genus and all other species in the dataset (inter-genus). The spread of scores are shown in Figure 1 and the minimum intra-genus and maximum inter-genus scores are outlined in Table 1. If the 94.5% boundary is used, we would expect no intra-genus score below 94.5% and no inter-genus score above 94.5% (Yarza *et al*., 2014). As expected, the spread of 16S intra-genus similarity scores were mostly above the 94.5% cut-off; however, both the original and new *Mycobacterium* showed a very small percentage of organisms that cross this threshold (<1%) (Table 1). Those scores below 94.5% tended to be between species such as *M. xenopi*, *M. heckeshornense*, and species at the edges of the phylogenetic divergence of the genera, such as *M. intracellulare* and *M. chelonae*, suggesting those species closely related to *M. xenopi* have more highly divergent 16S sequences than the rest of the mycobacteria. Unexpectedly, inter-genus similarity scores were often well above the 94.5% cut-off for all the genera, indicating general incongruences. Of all the genera, the original *Mycobacterium* had the lowest % of scores above 94% for inter-genus comparisons (Table 1).

**Figure 1.**
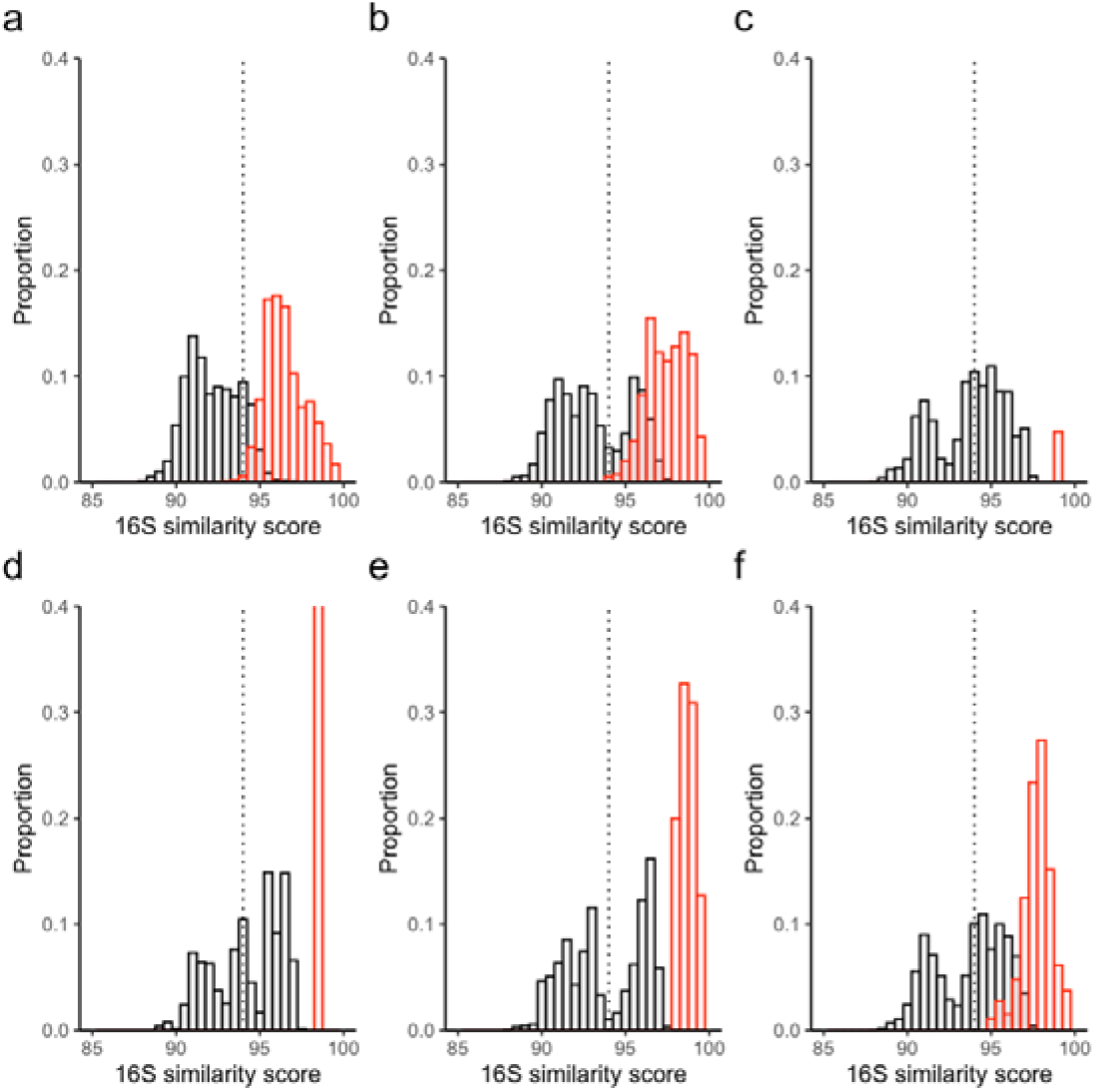
16S similarity scores between members of genera. Intra-genus scores are shown in red and inter-genus scores are shown in black. Histogram bin sizes are 0.5%. The proposed genus boundary of 94.5% is represented by the dashed line. Each genus is shown separately as follows: a) *Mycobacterium* (original) b) *Mycobacterium* (Gupta) c) *Mycobacteroides* d) *Mycolicibacillus* e) *Mycolicibacter* f) *Mycolicibacterium*. Note that the intra-genus proportion for *Mycolicibacillus* is 1.0 but the y-axis is cut at 0.4 for comparison between plots.

**Table 1.** 16S similarity scores. Inter-genus scores are those between members of the designated genus and all other species. Intra-genus scores are those only between members of the designated genus.

| Genus | Max. inter-genus 16S similarity (i.e., ideally $\leq 94.5\%$) | % of inter-genus scores above 94.5% | Min. intra-genus 16S similarity (i.e., ideally $> 94.5\%$) | % intra-genus scores below 94.5% |
| --- | --- | --- | --- | --- |
| *Mycobacterium* (original) | 97.57 | 16.61 | 92.73 | 0.5 |
| *Mycobacterium* (Gupta) | 97.78 | 35.95 | 93.83 | 0.09 |
| *Mycobacteroides* | 97.4 | 52.43 | 99.01 | 0 |
| *Mycolicibacillus* | 97.37 | 56.26 | 98.35 | 0 |
| *Mycolicibacter* | 97.78 | 46.82 | 97.85 | 0 |
| *Mycolicibacterium* | 97.77 | 53.78 | 94.95 | 0 |

Designation of the five newly proposed genera was originally based on phylogenetic grouping and AAI similarity within those groups (Gupta, Lo and Son, 2018). AAI scores above 65% are often observed for members of the same genus (Konstantinidis, Rosselló-Móra and Amann, 2017). Comparisons of inter- and intra-genus scores showed that for the original *Mycobacterium* genus, there is a clean split between these scores (Figure 2; Table 2). Thus, all non-mycobacteria genera had an AAI of <65% to any *Mycobacterium* (original), and all *Mycobacterium* (original) had an AAI score >65% to any other *Mycobacterium*. The intra-genus scores for all five new genera were also consistently above 64%. These combined results suggest that the split of Mycobacterium was not warranted.

**Figure 2.**
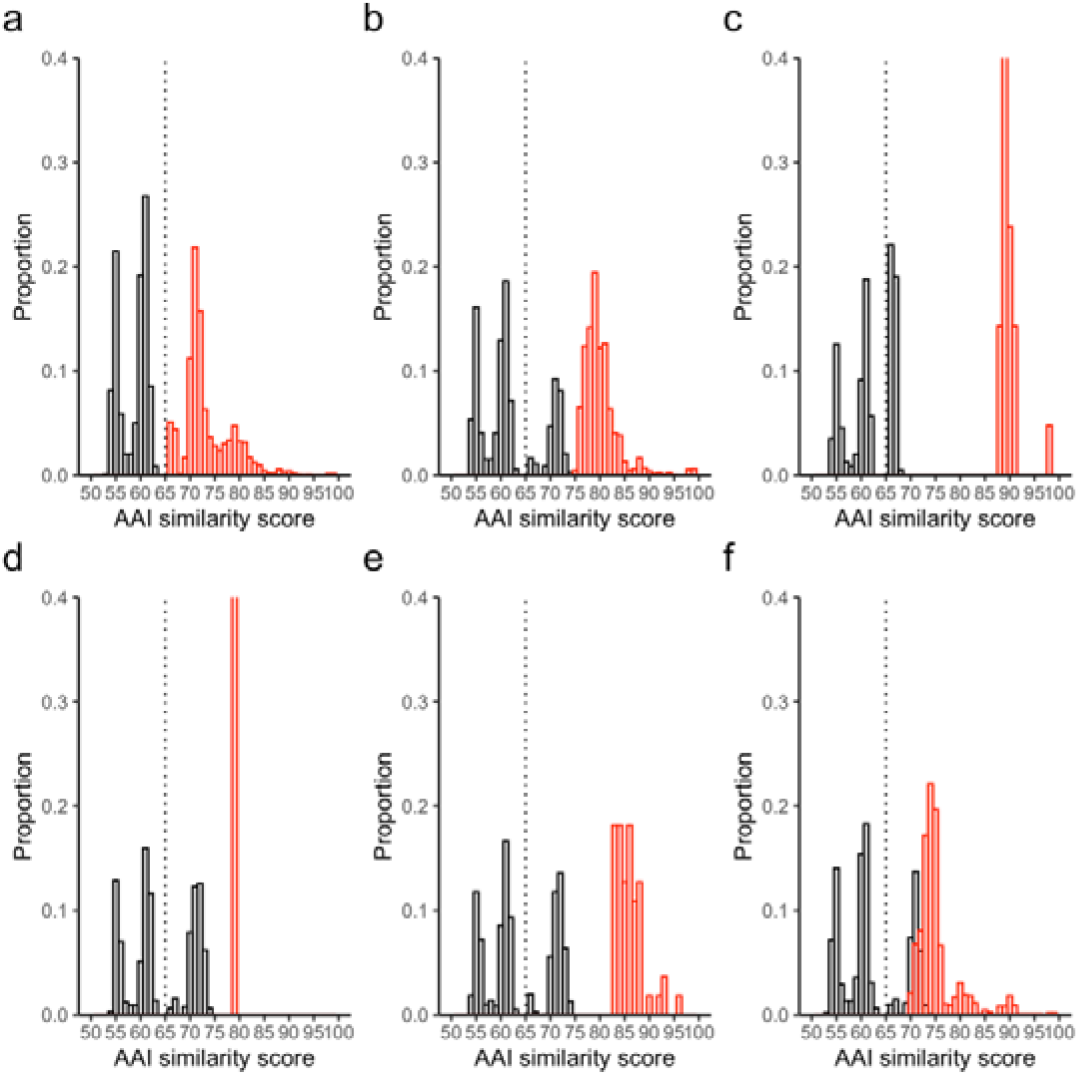
AAI similarity scores between members of genera. Intra-genus scores are shown in red and inter-genus scores are shown in black. Histogram bin sizes are 1%. The proposed genus AAI boundaries of 65% is represented by a dashed line. Each genus is shown separately as follows: a) *Mycobacterium* (original) b) *Mycobacterium* (Gupta) c) *Mycobacteroides* d) *Mycolicibacillus* e) *Mycolicibacter* f) *Mycolicibacterium*. Note that the intra-genus proportion for *Mycolicibacillus* is 1.0 but the y-axis is cut at 0.4 for comparison between plots.

**Table 2.** AAI similarity scores. Inter-genus scores are those between members of the designated genus and all other species. Intra-genus scores are those only between members of the designated genus.

| Genus | Max inter-genus AAI similarity (i.e., ideally $< 65\%$) | % inter-genus scores above 65% | Min intra-genus AAI similarity (i.e., ideally $> 65\%$) | % intra-genus scores below 65% |
| --- | --- | --- | --- | --- |
| *Mycobacterium* (original) | 63.75 | 0 | 65.48 | 0 |
| *Mycobacterium* (Gupta) | 74.37 | 28.02 | 74.85 | 0 |
| *Mycobacteroides* | 68.25 | 41.64 | 88.38 | 0 |
| *Mycolicibacillus* | 74.06 | 42.55 | 79.18 | 0 |
| *Mycolicibacter* | 74.37 | 40.9 | 83.1 | 0 |
| *Mycolicibacterium* | 73.91 | 31.99 | 69.83 | 0 |

### POCP supports original *Mycobacterium* genus over alternative sub-split

The POCP between members of the same genus (intra-genus) should be no less than 50% and the POCP between members of the genus and other species (inter-genus) should be no more than 50% (Qin *et al*., 2014). The original *Mycobacterium* genus fits this designation almost exactly, with only 0.031% of intra-genus POCP below 50% and 1.99% of inter-genus POCP above 50% (Figure 3; Table 3). The intra-genus scores of the new genera also all fit this criterion well. The maximum intergenus scores ranged from 55-88%, with 2-43% of comparisons above the 50% cut-off. The original *Mycobacterium* genus has 2% of the inter-genera comparisons >50% while the new *Mycobacterium* genus has a much higher percentage (25%) of the inter-genera comparisons above the 50% cut-off. The rest of the other genera had even higher proportions of the inter-genera comparisons >50% cut off. This indicates that the newly designated splits contain numerous species with relatively high identity to species from other genera.

**Figure 3.**
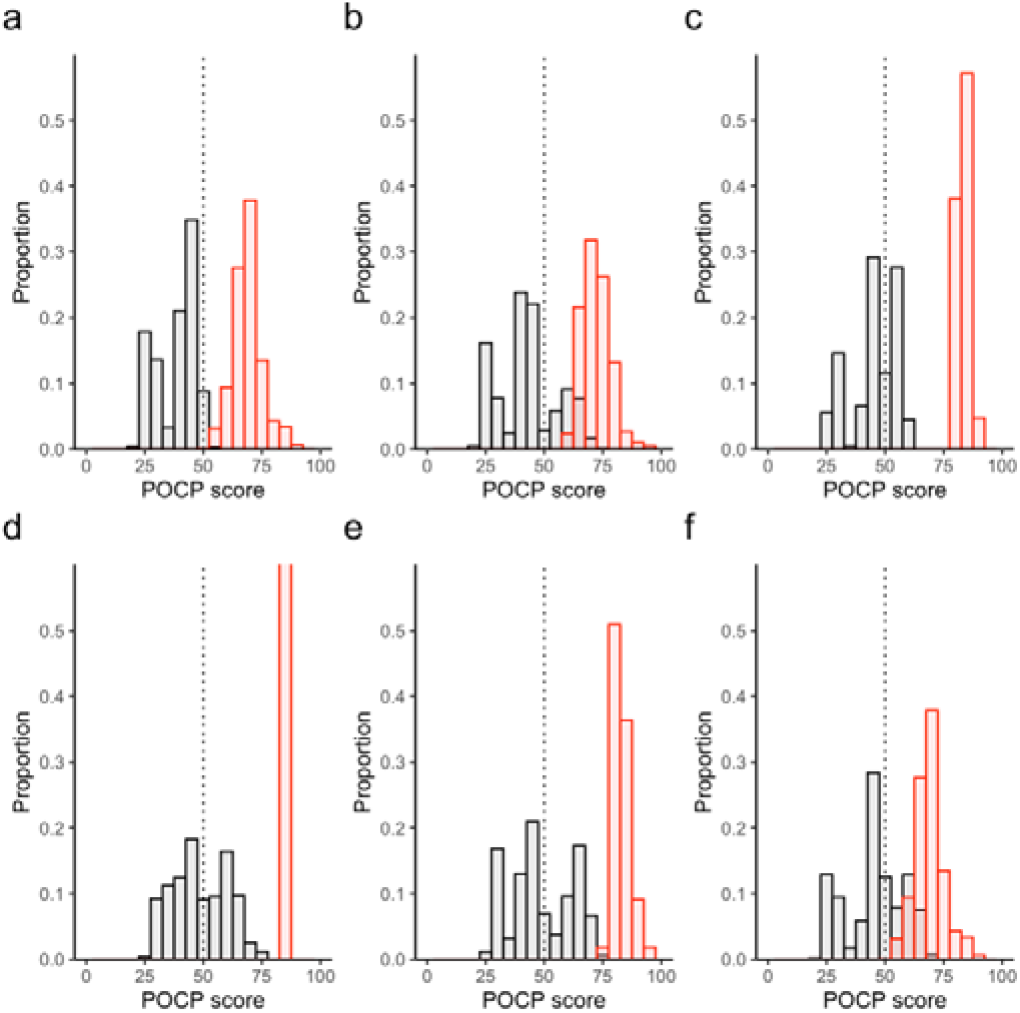
POCP scores between members of genera. Intra-genus scores are shown in red and inter-genus scores are shown in black. Histogram bin sizes are 5%. The proposed genus POCP boundary of 50% is represented by the dashed line. Each genus is shown separately as follows: a) *Mycobacterium* (original) b) *Mycobacterium* (Gupta) c) *Mycobacteroides* d) *Mycolicibacillus* e) *Mycolicibacter* f) *Mycolicibacterium*. Note that the intra-genus proportion for *Mycolicibacillus* is 1.0 but the y-axis is cut at 0.6 for comparison between plots.

**Table 3.** POCP scores. Inter-genus scores are those between members of the designated genus and all other species. Intra-genus scores are those only between members of the designated genus.

| Genus | Max inter-genus POCP similarity (i.e., ideally $< 50\%$) | % inter-genus scores above 50% | Min intra-genus POCP similarity (i.e., ideally $> 50\%$) | % intra-genus scores below 50% |
| --- | --- | --- | --- | --- |
| *Mycobacterium* (original) | 55.29 | 1.99 | 48.85 | 0.03 |
| *Mycobacterium* (Gupta) | 75.66 | 25.70 | 60.63 | 0 |
| *Mycobacteroides* | 87.87 | 41.91 | 79.32 | 0 |
| *Mycolicibacillus* | 75.23 | 43.23 | 83.35 | 0 |
| *Mycolicibacter* | 75.66 | 39.32 | 77.14 | 0 |
| *Mycolicibacterium* | 71.92 | 32.22 | 52.23 | 0 |

### Genus demarcation boundary supports the reconstitution of *Mycobacterium* with weak support for *Mycobacteroides*

Pairwise-comparisons against the type species of the proposed genus were made using all other species in that genus as well as other type species of closely-related genera with available genomes. These comparisons were used to determine the genus demarcation boundary to support/refute that grouping. A study by Barco *et al*. of over 850 genera found the mean alignment fraction (AF) and ANI to be 0.331 (95% CI: 0.308 −0.354) and 73.98% (95% CI: 73.34% −74.62%) respectively which defined the genus demarcation boundary (Barco et al., 2020). Thus, for a newly proposed genus, we would expect the AF and ANI of all species in that genus to be above these boundaries when compared to the proposed type species for that genus.

The comparison of all species within each proposed genus to the type species for that genus in terms of AF/ANI is outlined in Table 4 and Figure 4. The five newly proposed genera all had AF and ANI genus demarcation boundaries well above the 95% confidence interval of typical genera, indicating these new genera are too closely related to type species of other genera to be considered separate genera. Conversely, the original *Mycobacterium* genus had an ANI within the mean genus boundary confidence interval but an AF (0.265) considerably below the average genus demarcation boundary confidence interval (0.308 to 0.354). This supports its reconstitution as a genus over the proposed five-way split. However, the low AF value suggests that species at the genus boundary (i.e. those within the newly proposed *Mycobacteroides*) could represent a separate genus, but would need additional species in this clade to be sequenced to better support this two-way split.

**Figure 4.**
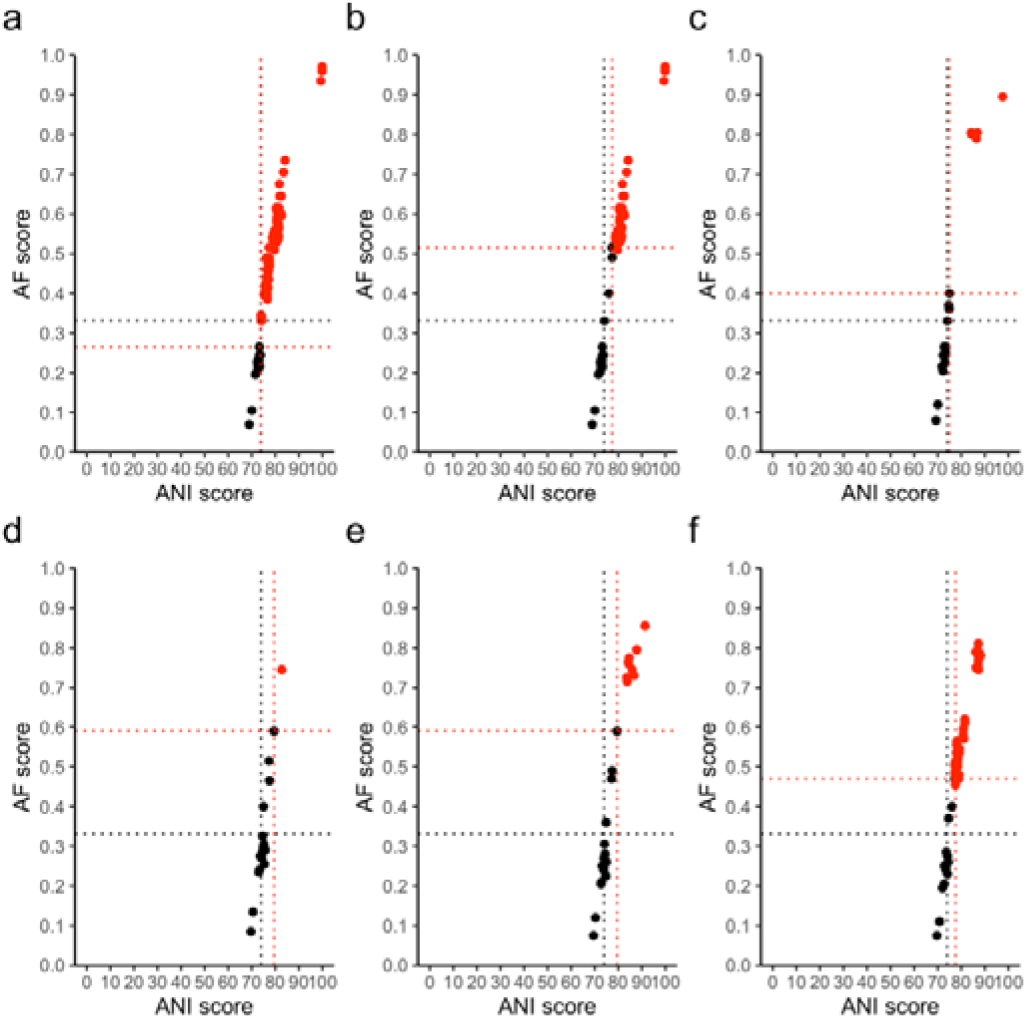
Alignment fraction (AF) and Average Nucleotide Identity (ANI) between genomes and the type species for each genus. Intra-genus scores are shown in red and inter-genus scores are shown in black. The proposed demarcation points for the genus is shown with a dashed red line. The mean demarcation points based on (Barco et al., 2020) are shown with dashed black lines. Each genus is shown separately as follows: a) *Mycobacterium* (original) b) *Mycobacterium* (Gupta) c) *Mycobacteroides* d) *Mycolicibacillus* e) *Mycolicibacter* f) *Mycolicibacterium*.

**Table 4.** Genus demarcation boundary. The AF and ANI scores of each proposed genus along with associated type species used are outlined. These values combined demarcate each individual genus boundary.

| Genus | Type species | AF boundary | ANI boundary |
| --- | --- | --- | --- |
| *Mycobacterium* (original) | *Mycobacterium tuberculosis* | 0.265 | 73.67 |
| *Mycobacterium* (Gupta) | *Mycobacterium tuberculosis* | 0.515 | 77.4 |
| *Mycobacteroides* | *Mycobacteroides abscessus* | 0.4 | 74.98 |
| *Mycolicibacillus* | *Mycolicibacillus trivialis* | 0.59 | 79.5 |
| *Mycolicibacter* | *Mycolicibacter terrae* | 0.59 | 79.49 |
| *Mycolicibacterium* | *Mycolicibacterium fortuitum* | 0.47 | 77.53 |

## Discussion

The taxonomic labelling of clinically important pathogens can have wide ranging implications in terms of correct, efficient identification and appropriate selection of treatment options. Of particular note, when renaming of species occurs, rule 56a of the Prokaryotic Code states that *nomen periculosum*, “a name whose application is likely to lead to accidents endangering health or life or both or of serious economic consequences” should be avoided (Parker, Tindall and Garrity, 2019). The genus *Mycobacterium* falls within this rule as it contains several strict and opportunistic pathogens. Thus, any renaming of species within this genus should be strongly supported before implemented.

A variety of genetic analyses were employed here to examine the evidence supporting the split of the genus *Mycobacterium* into five new genera, as proposed by (Gupta, Lo and Son, 2018). 16S rRNA similarity was found to be a very poor marker for delineation of the *Mycobacterium* genus, or new genera. The standard boundary of 94.5% similarity did not create clear separation between any of the genera proposed by Gupta *et al*. (Table 1, Figure 1). In each case, several species from one genus shared a 16S rRNA similarity above 94% with species from other genera. In the case of *Mycobacteroides*, *Mycolicibacillus* and *Mycolicibacterium* over 50% of the comparisons between species inside these genera and species designated outside were above this cut-off. Conversely, when comparing species inside these genera classifications with each other (e.g. *Mycobacteroides vs Mycobacteroides* or *Mycolicibacillus* vs *Mycolicibacillus*) the 94.5% boundary did classify species correctly, suggesting that the 16S boundary could be useful as an inclusionary boundary, but not exclusionary. However, for both the original and new *Mycobacterium* genera, 0.5% and 0.09% of comparisons were below this boundary, demonstrating that it is not completely clear cut.

Use of AAI scores was found to be better at demarcating genus boundaries. AAI clustering was used to define the new genera (Gupta, Lo and Son, 2018) and as expected, each genus formed groupings with high intra-genus AAI scores. However, as was seen with 16S, the inter-genera comparisons did not clearly separate species into the genera proposed by Gupta *et al*., with over 40% of inter-genus comparisons being below the 65% lower boundary cut-off for several of the newly proposed genera (Table 2, Figure 2). Conversely, all inter-genus comparisons for the original *Mycobacterium* genus were above the 65% boundary, strongly supporting this genus classification. Thus a sub-splitting is not warranted. The clustering and AAI scores more so suggest that the newly proposed genera are an expansion of the already defined complexes (Fedrizzi *et al*., 2017) and thus may represent sub-genus rankings instead.

The use of other genome-wide approaches for delineating genus boundaries further supports the re-amalgamation of the original *Mycobacterium* genus with the newly proposed genera forming sub-genus multi-species complexes. The POCP approach, using a score of shared protein coding genes across the genome, showed that the original *Mycobacterium* genus fell near the 50% proposed demarcation line (Table 3, Figure 3). Although the 50% boundary is somewhat arbitrary, the redefining of several pathogenic species into new genera is not supported by this genus demarcation approach and conversely supports the retainment of the original designation.

While the POCP utilises a non-standard approach, the (Barco *et al*., 2020) method uses AF and ANI, which is a standard genome relatedness index, to utilise genome similarity to look at genus boundary definitions. Comparing to the mean AF of 0.33 and ANI of 73.98% genus demarcation boundaries set by a large group of genus comparisons (Barco *et al*., 2020), the original *Mycobacterium* genus most closely approximates this boundary, with the five newly proposed genera being well above these points. Interestingly, *Mycobacteroides*, which contains the most basal species such as *M. abscessus* and *M. chelonae* also sits close to this boundary. Thus, these species could perhaps constitute the basis of a new genus in the future, but careful analysis of more genome sequences of closely related species would be needed to confirm this, especially in light of issues brought up by the analyses based on 16S rRNA gene, AAI, POCP, and ANI (e.g., inter-genera comparison results). Of note, due to the lack of *Mycobacterium* species with an ANI of 85-100%, the exact genus inflection points could not be determined, hence the use of genus demarcation instead. With the constant sequencing and discovery of new species, hopefully future analysis of AF and ANI scores can confirm these findings and determine the exact genus inflection point.

Overall, the variety of methods used here strongly support the reconstitution of the original *Mycobacterium* genus and the redesignation of the newly proposed genera as sub-genus complexes. Such complexes are useful clinically as many species within the same complex tend to share similar clinical and biochemical features. The conserved signature indels (CSIs) and conserved signature proteins (CSPs) for each of these groups as outlined by (Gupta, Lo and Son, 2018) can serve as markers for these complexes, as CSIs and CSPs can be used to denote any taxonomic rank or phylogenetic clade, not just a genus (Gao and Gupta, 2012; Gupta, 2019).

### Emended Description of the Genus *Mycobacterium* Lehmann and Neuman 1896 (Approved Lists 1980) (Skerman, McGowan and Sneath, 1980)

Mycobacterium (My.co.bac.te ri.um. Gr. n. mykes a fungus; N.L. neut. n. bacterium, a small rod; N.L. neut. n. Mycobacterium, a fungus rodlet).

The type species is *Mycobacterium tuberculosis* (Zopf, 1883) Lehmann and Neumann 1896 (Approved Lists 1980) (Skerman, McGowan and Sneath, 1980).

The characteristics of this genus match those of the emended description of the genus *Mycobacterium* as outlined in (Gupta, Lo and Son, 2018). All species in the genera *Mycobacterium, Mycobacteroides, Mycolicibacillus, Mycolicibacter* and *Mycolicibacterium* are included in this genus. They can be separated from other species in the family *Mycobacteriaceae* using both the signature CSI and CSP for these groups outlined in (Gupta, Lo and Son, 2018) as well as having an AF above 0.265 and ANI above 73.76% when compared to the type species, *Mycobacterium tuberculosis*.

## Acknowledgements

The authors thank Oren Tzfadia for his useful comments on the manuscript.

## References

Adeolu, M. et al. (2016)’Genome-based phylogeny and taxonomy of the “Enterobacteriales”: Proposal for enterobacterales ord. nov. divided into the families Enterobacteriaceae, Erwiniaceae fam. nov., Pectobacteriaceae fam. nov., Yersiniaceae fam. nov., Hafniaceae fam. nov., Morganellaceae fam. nov., and Budviciaceae fam. nov’, International Journal of Systematic and Evolutionary Microbiology. Microbiology Society, 66(12), pp. 5575–5599. doi: 10.1099/ijsem.0.001485.

Altschul, S. F. et al. (1990) ‘Basic local alignment search tool’, J Mol Biol, 215(3), pp. 403–410. doi: 10.1016/S0022-2836(05)80360-2.

Barco, R. A. et al. (2020) ‘A genus definition for bacteria and archaea based on a standard genome relatedness index’, mBio. American Society for Microbiology, 11(1). doi: 10.1128/MBIO.02475-19.

Fedrizzi, T. et al. (2017) ‘Genomic characterization of Nontuberculous Mycobacteria’, Scientific Reports. Nature Publishing Group, 7(1), p. 45258. doi: 10.1038/srep45258.

Gagneux, S. (2018) ‘Ecology and evolution of Mycobacterium tuberculosis’, Nature Reviews Microbiology, 16(4), pp. 202–213. doi: 10.1038/nrmicro.2018.8.

Gao, B. and Gupta, R. S. (2012) ‘Phylogenetic Framework and Molecular Signatures for the Main Clades of the Phylum Actinobacteria’, Microbiology and Molecular Biology Reviews. American Society for Microbiology, 76(1), pp. 66–112. doi: 10.1128/mmbr.05011-11.

GTDB - Mycobacterium tree (2021). Available at: https://gtdb.ecogenomic.org/tree?r=g_Mycobacterium (Accessed: 9 February 2021).

Gupta, A. K. (2015) ‘Classification’, in Springer Geology. Springer, pp. 69–87. doi: 10.1007/978-81-322-2083-1_3.

Gupta, R. S. (2019) ‘Commentary: Genome-Based Taxonomic Classification of the Phylum Actinobacteria’, Frontiers in Microbiology. Frontiers Media S.A., 10(FEB), p. 206. doi: 10.3389/fmicb.2019.00206.

Gupta, R. S., Lo, B. and Son, J. (2018) ‘Phylogenomics and Comparative Genomic Studies Robustly Support Division of the Genus Mycobacterium into an Emended Genus Mycobacterium and Four Novel Genera’, Frontiers in Microbiology. Frontiers, 9, p. 67. doi: 10.3389/fmicb.2018.00067.

Gupta, R. S., Lo, B. and Son, J. (2019) ‘Corrigendum: Phylogenomics and Comparative Genomic Studies Robustly Support Division of the Genus Mycobacterium into an Emended Genus Mycobacterium and Four Novel Genera’, Frontiers in Microbiology. Frontiers Media S.A., 10(APR), p. 714. doi: 10.3389/fmicb.2019.00714.

Konstantinidis, K. T., Rosselló-Móra, R. and Amann, R. (2017) ‘Uncultivated microbes in need of their own taxonomy’, ISME Journal. Nature Publishing Group, pp. 2399–2406. doi: 10.1038/ismej.2017.113.

Konstantinidis, K. T. and Tiedje, J. M. (2005) ‘Towards a genome-based taxonomy for prokaryotes’, Journal of Bacteriology. American Society for Microbiology Journals, 187(18), pp. 6258–6264. doi: 10.1128/JB.187.18.6258-6264.2005.

Lehmann, K. B. and Neumann, R. (1896) Atlas und Grundriss der Bakeriologie und Lehrbuch der speziellen bakteriologischen Diagnositk. 1st editio.

Lopes-Santos, L. et al. (2017) ‘Reassessment of the taxonomic position of Burkholderia andropogonis and description of Robbsia andropogonis gen. nov., comb. nov’, Antonie van Leeuwenhoek, International Journal of General and Molecular Microbiology. Springer Netherlands, 110(6), pp. 727–736. doi: 10.1007/s10482-017-0842-6.

Nicholson, A. C. et al. (2018) ‘Revisiting the taxonomy of the genus Elizabethkingia using whole genome sequencing, optical mapping, and MALDI-TOF, along with proposal of three novel Elizabethkingia species: Elizabethkingia bruuniana sp. nov., Elizabethkingia ursingii sp. nov., and Elizabethkingia occulta sp. nov.’, Antonie van Leeuwenhoek, International Journal of General and Molecular Microbiology. Springer Netherlands, 111(1), pp. 55–72. doi: 10.1007/s10482-017-0926-3.

Parker, C. T., Tindall, B. J. and Garrity, G. M. (2019) ‘International code of nomenclature of Prokaryotes’, International Journal of Systematic and Evolutionary Microbiology. Microbiology Society, 69(1), p. S1. doi: 10.1099/ijsem.0.000778.

Parks, D. H. (2014) CompareM. Available at: https://github.com/dparks1134/CompareM.

Parks, D. H. et al. (2015) ‘CheckM: assessing the quality of microbial genomes recovered from isolates, single cells, and metagenomes.’, Genome research, p. gr.186072.114–. doi: 10.1101/gr.186072.114.

Parks, D. H. et al. (2018) ‘A standardized bacterial taxonomy based on genome phylogeny substantially revises the tree of life’, Nature Biotechnology. Nature Publishing Group, 36(10), p. 996. doi: 10.1038/nbt.4229.

Parks, D. H. et al. (2020) ‘A complete domain-to-species taxonomy for Bacteria and Archaea’, Nature Biotechnology.Nature Research, pp. 1–8. doi: 10.1038/s41587-020-0501-8.

Qin, Q. L. et al. (2014) ‘A proposed genus boundary for the prokaryotes based on genomic insights’, Journal of Bacteriology. American Society for Microbiology, 196(12), pp. 2210–2215. doi: 10.1128/JB.01688-14.

R Core Team (2017) ‘R: A language and environment for statistical computing.’ R Foundation for Statistical Computing, Vienna, Austria. Available at: http://www.r-project.org/.

Seemann, T. (2014) ‘Prokka: rapid prokaryotic genome annotation’, Bioinformatics, 30(14), pp. 2068–2069. doi: 10.1093/bioinformatics/btu153.

Skerman, V. B. D., McGowan, V. and Sneath, P. H. A. (1980) ‘Approved lists of bacterial names’, International Journal of Systematic Bacteriology. Microbiology Society, 30(1), pp. 225–420. doi: 10.1099/00207713-30-1-225.

Tortoli, E. et al. (2017) ‘The new phylogeny of the genus Mycobacterium: The old and the news’, Infection, Genetics and Evolution. Elsevier, 56, pp. 19–25. doi: 10.1016/J.MEEGID.2017.10.013.

Tortoli, E. et al. (2019) ‘Genome-based taxonomic revision detects a number of synonymous taxa in the genus Mycobacterium’, Infection, Genetics and Evolution. Elsevier, p. 103983. doi: 10.1016/J.MEEGID.2019.103983.

Warnes, G. R., Bolker, B. and Lumley, T. (2012) gplots: Various R programming tools for plotting data. R package version 2.6.0. Available at: http://cran.r-project.org/web/packages/gplots/.

Yarza, P. et al. (2008) ‘The All-Species Living Tree project: A 16S rRNA-based phylogenetic tree of all sequenced type strains’, Systematic and Applied Microbiology. Syst Appl Microbiol, 31(4), pp. 241–250. doi: 10.1016/j.syapm.2008.07.001.

Yarza, P. et al. (2014) ‘Uniting the classification of cultured and uncultured bacteria and archaea using 16S rRNA gene sequences’, Nature Reviews Microbiology. Nature Publishing Group, 12(9), pp. 635–645. doi: 10.1038/nrmicro3330.

Zopf, W. (1883) Die Spaltpilze.□: Nachdem neuesten Standpunkte. Breslau: Eduard Trewendt.

